# RNA editing in inflorescences of wild grapevine unveils association to sex and development

**DOI:** 10.1101/2020.03.03.974626

**Authors:** Miguel J. N. Ramos, David Faísca-Silva, João L. Coito, Jorge Cunha, Helena Gomes Silva, Wanda Viegas, M. Manuela R. Costa, Sara Amâncio, Margarida Rocheta

**Affiliations:** Linking Landscape, Environment, Agriculture and Food (LEAF), School of Agriculture, University of Lisbon, Tapada da Ajuda, 1359-017 Lisboa, Portugal; Instituto Nacional de Investigação Agrária e Veterinária, Quinta d’Almoinha, Dois Portos, Portugal; Biosystems and Integrative Sciences Institute (BioISI), Plant Functional Biology Centre, University of Minho, Campus de Gualtar, 4710-057 Braga, Portugal

**Keywords:** *Vitis vinifera sylvestris*, RNA editing, development, nuclear transcripts, dioecy, inflorescence, sex

## Abstract

RNA editing challenges the central dogma of molecular biology, by modifying the genetic information at the transcription level. Recent reports, suggesting increased levels of RNA editing in plants, raised questions on the nature and dynamics of such events during development. We here report the occurrence of distinct RNA editing patterns in wild *Vitis* flowers during development, with twelve possible RNA editing modifications observed for the first time in plants. RNA editing events are gender and developmental stage specific, identical in subsequent years of this perennial species and with distinct nucleotide frequencies neighboring editing sites on the 5’ and 3’ flanks. The transcriptome dynamics unveils a new regulatory layer responsible for gender plasticity enhancement or underling dioecy evolution in *Vitis*.

## INTRODUCTION

It is well consensual that DNA provides the code for synthesizing RNA and proteins, however, there is not always a direct one-to-one relationship between DNA, RNA, and protein. Co-transcriptional and post-transcriptional processing, such as RNA splicing and RNA editing, can alter the structure and sequence of RNAs, leading to different transcripts produced from the same DNA sequence. RNA editing can be defined as any site-specific alteration in RNA sequences, that include insertion or deletion of nucleotides and base conversion, and is an effective mechanism to modifying and enrich genetic information beyond the genetic code (Bahn et al., 2012; Li et al., 2011; Liscovitch-Brauer et al., 2017; Qulsum et al., 2019). The predominant type of RNA editing in animals is the conversion of adenosine (A) to inosine (I), catalyzed by a family of adenosine deaminases (ADARs) acting on double strand RNA (Nishikura, 2006). This editing is also known as A-to-G editing since inosine in RNA is interpreted as guanosine (G) by the translational machinery (Nishikura, 2006). On the other hand, ADAR-like enzymes were not yet found in plants (Jin et al., 2009). Another well documented type of RNA editing in humans is the deamination of cytidine leading to uridine (C-to-U), a RNA editing event catalyzed by the activation of APOBECs (Salter et al., 2016). However, it is still an enigma how the remaining ten nucleotide modifications found in humans (Li et al., 2011) and in cephalopods (Liscovitch-Brauer et al., 2017) transcriptomes can occur.

In the plant kingdom, RNA editing is widespread in organellar transcripts of almost all land plants, mainly involving the deamination of cytidines into uridines (C-to-U) by specific pentatricopeptide repeat (PPR) proteins, encoded in the nuclear genome (Takenaka et al., 2019). There are two main classes of PPR proteins: P and PLS (Ichinose and Sugita, 2017). PLS seems to be the most relevant class on plant RNA editing (Rudinger et al., 2008) due to a conserved extended domain (E) at the C terminus (PPR-E) (Ichinose and Sugita, 2017; Okuda et al., 2009; Okuda et al., 2007), which contains a DYW motif (Ichinose and Sugita, 2017). The DYW motif encodes a cytidine deaminase active center, probably responsible for C-to-U RNA editing (Rudinger et al., 2008). Although, the specificity of PPR-DYW is yet undetermined, PPRs are in general considered as non-target specific. Reverse U-to-C editing was observed in hornworts, lycophytes and ferns (Knie et al., 2016) and seems to be rare in higher plants. A global vision of RNA editing in plant nuclear transcripts was described in *Arabidopsis* (Meng et al., 2010) and recently discriminated between different plant tissues (Qulsum et al., 2019).

Although the previous studies shed some light in plant editing, several important questions related to mRNA modifications in plants still need to be answered, namely: (1) Is RNA editing related to sex in dioecious plants? (2) Are RNA editing patterns similar in successive years in perennial plants? (3) Is editing transient or permanent during flower development? Finally, (4) are ADAR or APOBEC-like sequences present and expressed in plant genomes?

*Vitis* is a perennial genus that in European countries has only one growing season, when it reiterates the development program allowing the analysis of the same plant on subsequent years. The present work in dioecious *Vitis vinifera sylvestris* (hereafter referred as *V. v. sylvestris*) lies in the difference between the two sexes: abortion of carpel development in male flowers and reflex stamens with non-viable pollen in female plants (Badouin et al., 2020; Coito et al., 2017; Massonnet et al., 2020; Picq et al., 2014; Zhou et al., 2017). We compared RNA edition events at several specific developmental stages, disclosing for the first time in plants, the twelve nucleotides modifications, previously detected in edited transcripts from human cells and cephalopods (Li et al., 2011; Liscovitch-Brauer et al., 2017). We found that RNA editing events have specific profiles in consecutive years, developmental stages of inflorescences and distinct sexes. The discovery of transcripts from adenosine and cytidine deaminase-like sequences, during inflorescence development, is moreover a marked breakthrough for a deeper understanding of RNA editing in plants. These findings suggest that RNA editing is an essential highly controlled mechanism allowing the quick synthesis of all protein isoforms without resorting to DNA modifications. Even very highly distributed, the complete mechanism of RNA editing still need to be discovered.

## RESULTS

### Pipeline to identify and validate RNA-DNA Differences

We started by re-sequencing the genomes of male and female inflorescences at different developmental stages (B, D and H) (Baggiolini, 1952), from the same individuals whose transcriptomes were previously sequenced (Ramos et al., 2014) (Figure 1). The sequencings were performed using an Illumina pipeline and reads length ranged between 50 bp for RNA-seq and 2 x 125 bp for genomes re-sequencing. Reads were inspected to remove low quality calls and adapters, as detailed in **Methods** section. Good quality reads were mapped against a reference genome, sequenced from a hermaphrodite variety, derived from Pinot Noir (*Vitis vinifera vinifera* cv. PN40024 12X) (Jaillon et al., 2007; Vitulo et al., 2014). Transcripts abundance estimation was assessed by CLC Genomics Workbench, considering the number of Reads per Kilobase of exon model per Million mapped reads (RPKMs) (Ramos et al., 2014).

**Figure 1.**
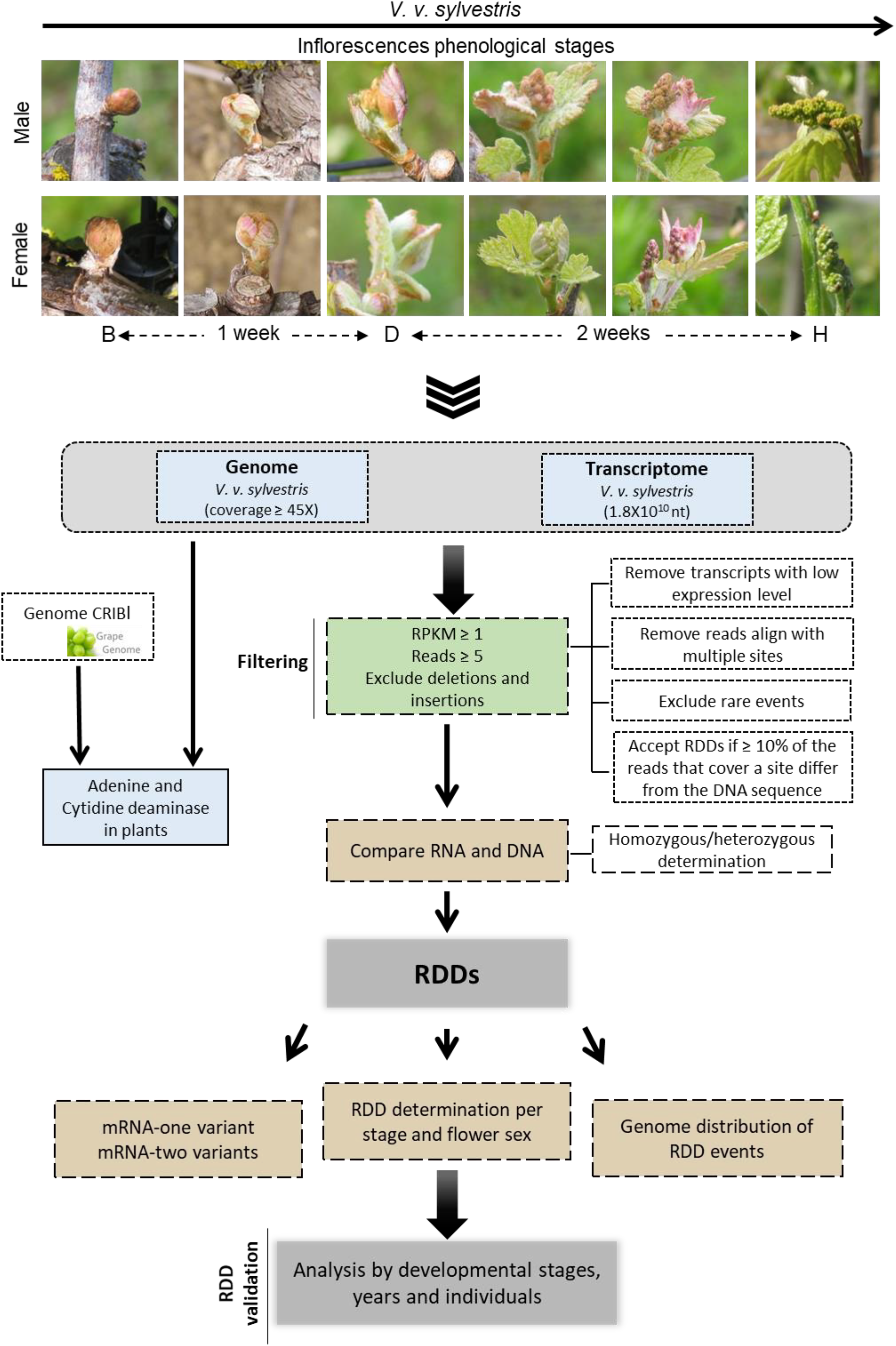
Pipeline to identify and validate RNA-DNA differences. To identify RDDs, both genome and transcriptomes were compared at three development stages of male and female *Vitis vinifera sylvestris*. Several filters were considered: 1) RPKMs must be equal or higher than one in all developmental stages in each flower type; 2) reads must be equal or higher than five; 3) deletions, insertions or indels were excluded. Results were validated on several inflorescence samples collected from the same male and female plants in the same time-frame corresponding to the phenological stages B, D and H (Baggiolini, M. (1952)). B samples were collected at the beginning of floral bud; D samples were collected one week later; H samples were collected two weeks from D.

Nucleotide modifications search was performed using re-sequenced genomes and transcriptomes of the same individuals, both assembled against the fully annotated reference genome (see **Methods** for more details) (Figure 1). Differences between *V. v. sylvestris* re-sequenced genomes and transcriptomes were detected through the identification of single nucleotide modifications (see **Methods** for more details). Shortly, sequences from RNA-seq and genomes re-sequencing were compared using the following four major filters (Figure 1): first, only genes with simultaneous expression in the six samples analyzed (17356) (RPKM ≥ 1) were considered, allowing the exclusion of false negatives; second, considering the low error rate of Illumina short read sequencing (0.001) (Glenn, 2011; Schirmer et al., 2016), we imposed that all analyzed *loci* should have five or more reads, corresponding to a cumulative error rate of 0.001^5^; third, since all reads that aligned against multiple sites could lead to mismapping errors, repetitive regions and Insertions/Deletions (InDels) were removed to reduce analysis complexity; and four, nucleotides differences observed in less than 10% of the reads were discarded (Figure 1).

In this manuscript, the designations previously used in human editing changes (Li et al., 2011) are here adopted, so that the occurrence of a difference between RNA and DNA sequences (RDDs) is referred as an "event", and the chromosomal location where it occurs as the "site".

To validate the *in silico* results, we collected samples from the same plants at similar flower developmental stages used in the original sequencing (Ma1_2014 and Fe1_2014), at subsequent years. Additionally, samples from other male and female plants were also collected in different years. These samples allowed a very large characterization of RNA editing events in specific flower developmental stages, in several individuals of the same sex over the years.

### Dynamics of RDDs in *V. v. sylvestris* during inflorescence development

To access the dynamics of RNA editing levels during inflorescence development, we analyzed six *V. v. sylvestris* transcriptome samples, three from male and three from female plants. Although the full *Vitis* genome is available (Jaillon et al., 2007) it could not be used to establish a precise comparison between DNA-RNA, since the subject of the present research concerns the wild dioecious subspecies *V. v. sylvestris*. Thus, to accurately determine editing sites in both male (Ma1_2014) and female (Fe1_2014) plants, the genomes of these plants were re-sequenced (Ramos et al., 2020).

Considering only the 17356 genes with expression (RPKM ≥ 1) in the three stages of flower development from male and female plants, we evaluated the number of genes showing RNA editing events. A difference in the number of edited RNAs between male and female flowers across development was observed (Figure 2A). Male flowers consistently showed similar number of genes with edited transcripts at the three developmental stages (2678, 2735 and 2801, for stages B, D and H, respectively), contrasting with female flowers, that had a very high amount of edited transcripts at stages D and H (10955 and 10458, respectively) compared with the lower level observed at stage B (2238) (Figures 2A and S1). The results showed that only 23.2% of edited transcripts were common to both sexes being 72.7% female specific (Figure 2B). The analysis of RNA editing events, by developmental stages, also disclosed a marked disparity between male and female inflorescences, as male flowers show a more stable behavior in terms of RNA editing (Figure 2B). Moreover, editing/gene frequencies were 2.5 and 6.3 in male and female inflorescences, respectively, which was accompanied by a higher number of female exclusive RDDs events (83292) (Figures 2C and S1).

**Figure 2.**
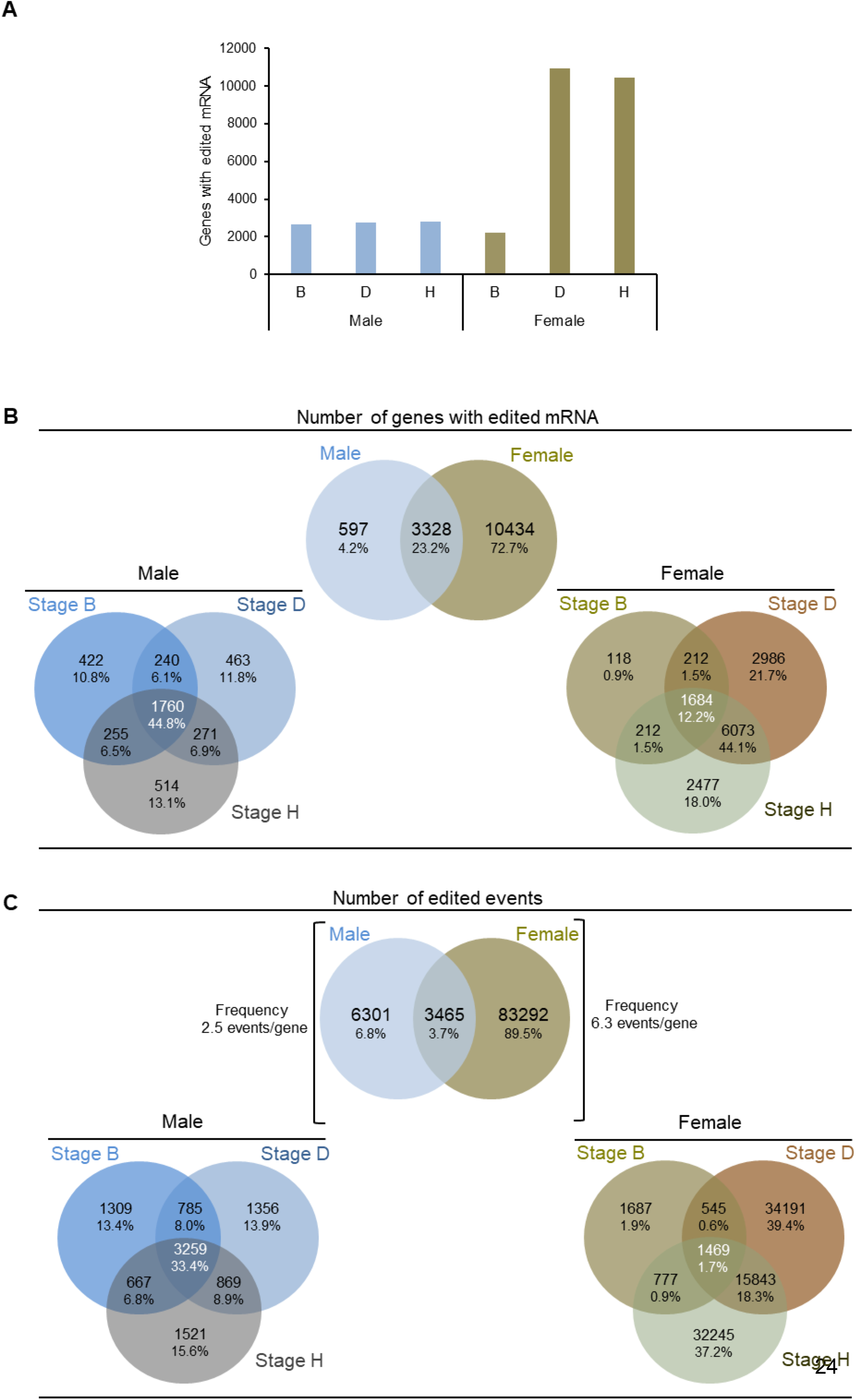
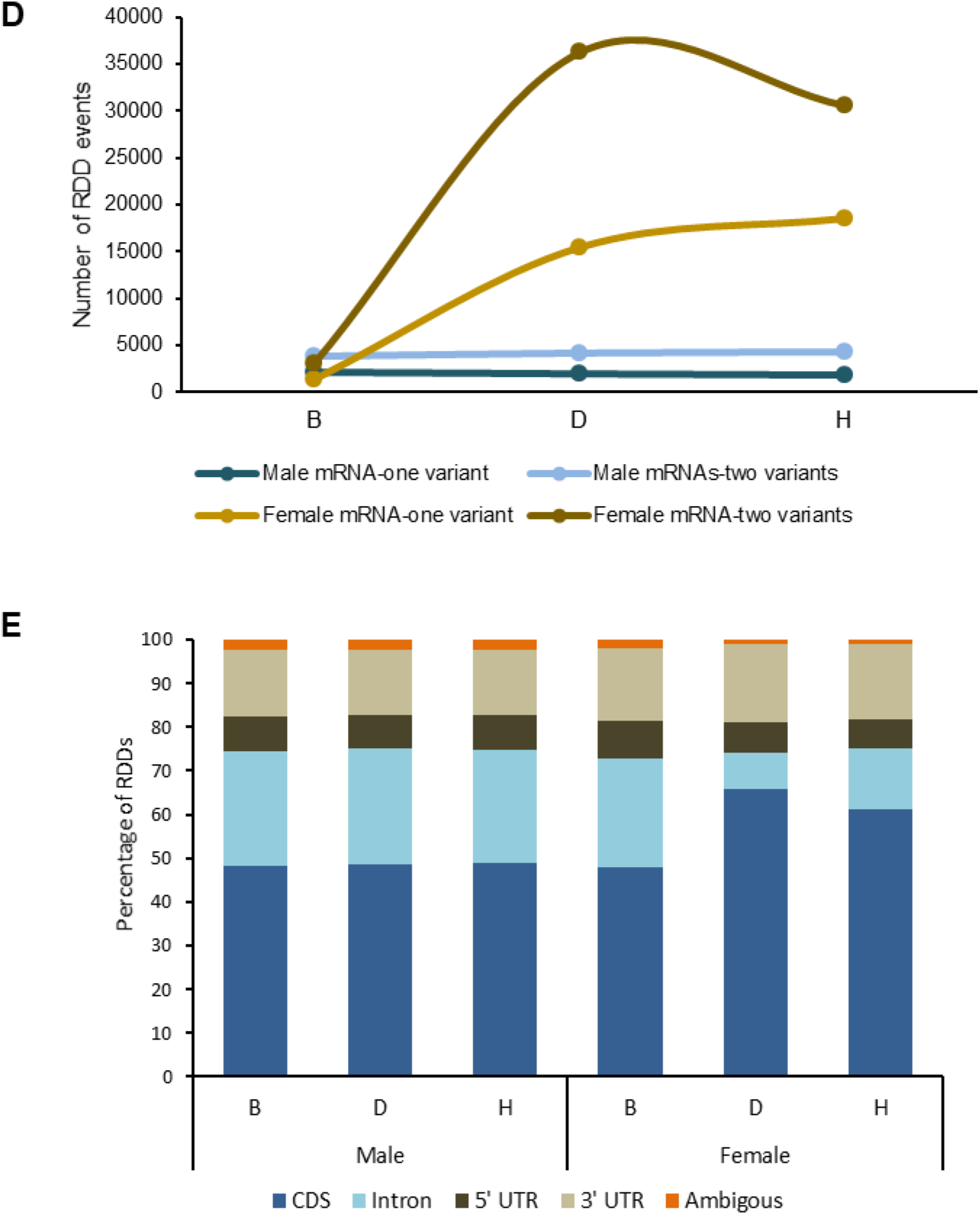
Dynamic of RDD modifications in *V. v. sylvestris* during flower development. (A) There is an increasing number of genes with edited mRNAs in stages D and H of female flower development (to the percentage of modifications in each stage see Figure S1). The same is not observed in male samples, where the number of editing sites remains in the same order of magnitude. Filters used: reads ≥ 5 and RPKM ≥1. (B) The number of transcripts with nucleotide modifications between DNA and RNA, compared in male and female samples (upper diagram) showed a large number of edited mRNA transcripts in female inflorescence. Regarding flower development, by flower type (stages B, D and H – lower diagrams), it was possible to observe a large percentage of edited transcripts common to all development stages in male plant. This observation contrasts with female flower where only 12.2% of edited transcripts are common to the three development stages. Filters used: reads ≥ 5 and RPKM ≥1. (C)The editing frequency, corresponding to the average number of RDD events per transcript (upper diagram), is higher in the female plant than in male individuals and only 3.7% of the sites are shared between flower types. The lower diagrams show an increasing percentage of editing events in the later development stages of female flower due to exclusive RDD sites. Filters used: reads ≥ 5 and RPKM ≥1. (D) For each edited RNA transcript one of two situations was observed: 1) only the edited mRNA was identified (mRNA-one variant) or 2) both the edited and non-edited mRNAs were identified (mRNA-two variants). The number of RDD events producing one mRNA (mRNA-one variant) or two distinct mRNAs (mRNA-two variants) in male plant seems to be stable during development. Female individual, in contrast, showed an increasing number of RDDs events in D and H development stages. In both flower types the number of mRNA-two variants is higher, indicating that RDD can occur only in some mRNA molecules from the same gene. (E) The increased percentage of RDDs in female coding region (CDS), stages D and H, is collinear with the decrease of intronic RDDs. Additionally, there is a higher number of RDD events in the 3’ UTR than in the 5’ UTR. Filters used: reads ≥ 5 and RPKM ≥1.

In-depth analysis revealed that male inflorescences are heterozygous in 33% of female exclusive events (27716), contrasting with male specific RNA editing events (6301) where the female inflorescences are heterozygous in 8% (483) (Ramos et al., 2020). By being heterozygous, male flowers do not need RNA editing to translate the same proteins as female flowers.

Besides evidences of RNA editing at homozygous and heterozygous gDNA sites, the subsequent analyzes were restricted to homozygous sites to reduce complexity. Considering the possibility of a gene having two mRNA variants (edited and unedited), the dynamics of both forms throughout inflorescence development was also analyzed. At this point, for clarity, when all transcripts presented only the edited RNA form will here be referred as mRNA-one variant; and when transcripts simultaneously present edited and unedited forms will be referred as mRNA-two variants. Again, male and female flowers had distinct profiles during development (Figure 2D). While male inflorescences displayed similar levels of both mRNA variants, female inflorescences showed a significant increase in RDD events in stage D, associated with a marked discrepancy between mRNA-one and mRNA-two variants (Figure 2D).

Taking into consideration that in both sexes all genomic regions were affected by editing, the observed increase in editing events in coding regions (CDS) during intermediate and later developmental stages of female inflorescences (Figure 2E), suggests a potential need for more protein isoforms in addition to those encoded by DNA. Although, it may be argued that mRNA-edited transcripts will not necessarily be translated into proteins, it must be emphasized that previous proteomic validation in human cells proved the existence of new proteins different from the ones encoded by the respective genomic DNA sequences (Li et al., 2011; Maas et al., 2001).

The present study shows that from an editing point of view, the development of male inflorescences seems to be less demanding than the female ones. This differential RNA editing profiles may be in part explained by the heterozygous levels of both flowers types. RDD patterns are associated to distinct developmental stages of male and female flowers, probably underling dioecy evolution in *Vitis*.

### Twelve recoding types are detected in *Vitis*

Our data revealed all the 12 possible mismatches between DNA and RNA across B, D and H inflorescence development stages, in wild grapevine, as already found in human (Li et al., 2011) and cephalopod (Liscovitch-Brauer et al., 2017). Through the use of four major filters (see **Methods**) and considering only homozygous sites, male flowers showed a higher number of nucleotide modifications during inflorescence development (Figure 3A), contrasting with the stage B of female flowers with the lowest number of editing events across all modifications (Figure 3B). The normalization by percentage (number of modifications over total number of RDDs at each developmental stage) of male and female RDDs numbers revealed that in stage B of female inflorescence, the number of transitions were lower when compared with transversions (Figure S2).

**Figure 3.**
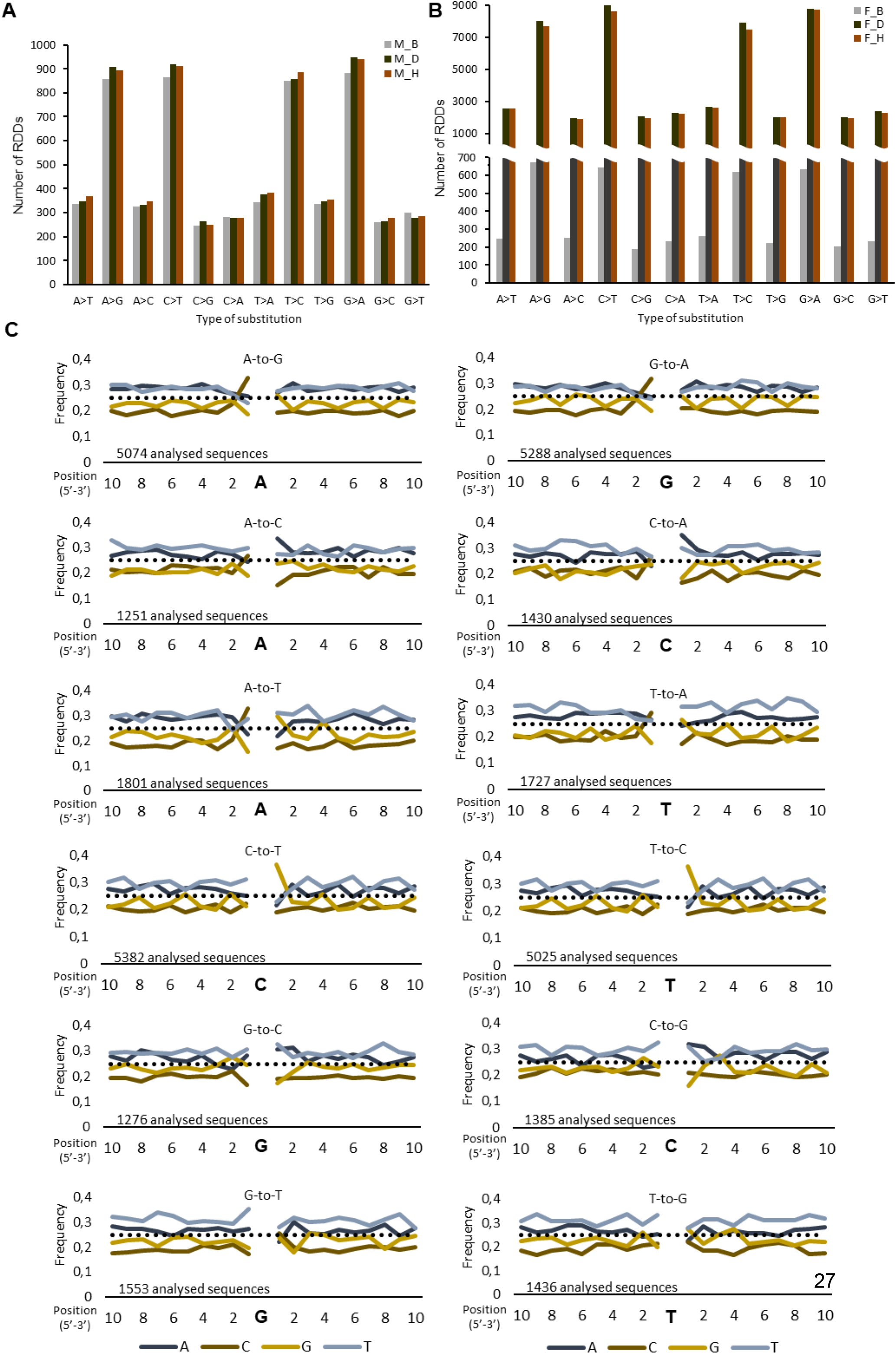
Twelve recoding types are detected in *Vitis*. (A and B) The transcriptome of *V. v. sylvestris* male (A) and female (B) inflorescences in three developmental stages (B, D and H) reveals twelve possible modification types, with purines to purines and pyrimidines to pyrimidines showing higher values. See **Figure S2**. (C) To access the nucleotides around the edited sites, both female and male genomes were joined together and sequences were analyzed from 100 nt upstream, to 100 nt downstream of the modification site, but we only show the 10 nt from each site. Whenever the RDD event involves an adenine (six first plots) there is enrichment in cytosine at the 5’ site. At the 3’side enrichment in purine bases seems favored with the exception of A-to-G and G-to-A changes, where no enrichment is observed, and T-to-G and G-to-T, where, again, enrichment in cytosine is observed. The horizontal line at a frequency at 0.25 indicates the expected frequency if the four nucleotides are randomly represented (see Table S1).

Recently, 9 of these nucleotide changes were also reported in seedlings, leaves and stem of *Arabidopsis* tissues (Qulsum et al., 2019). The *Arabidopsis* unreported C-to-G, C-to-A and G-to-T modifications were found in *Vitis* inflorescences with patterns similar to the remaining transversions (Figures 3A and 3B; Figure S2).

Since only a partial explanation for these modifications is available in plants (Cheng et al., 2016), we searched for other potential patterns in regions surrounding edited sites of all RDD types (Figure 3C). Comparisons between the genomic features and the six transcriptomes were analyzed at 1000 nt upstream and downstream of edited site, applying a Tukey’s fences method (k = 1.5), to determine if the dataset displayed outliers or extreme values (see Table S1). The data revealed different sequence contexts at 5’ and 3’ sides surrounding the RDD nucleotide (Figure 3C). First, when adenine (A) was involved, the base 5’ adjacent to the editing sites contained an enrichment in cytosine (C). The base 3’ however, seemed to be more permissive showing an enrichment in one of the purine bases (A or G) with four exceptions: A-to-G, G-to-A and G-to-C where no enrichment or depletion occurred, and G-to-T where simultaneously occurred an enrichment in C (Figure 3C and Table S1). Also, the base 5’ adjacent to adenosine in A-to-G sites was depleted in guanosine (G) while the base 3’ was enriched in G (Figure 3C and Table S1), a result consistent with previous studies in humans (Wang et al., 2013). Additionally, the motif discovery tool MEME Suite (Bailey et al., 2009) was used to search motifs for RNA-binding proteins in edited transcripts, but although we have performed our analysis on 100 nt upstream and downstream of edited sites, results were inconclusive.

### Genomic regions and protein synthesis are differentially affected by RDDs

We combined the analysis of both genomes to study the distribution of the twelve types of RNA recoding events on the various genomic regions, revealing the following patterns: (1) the RDDs in CDS regions ranged between 53% and 67%; (2) the modifications in introns varied between 12% and 16%; (3) there was a preference for editing at the 3’ UTR (14% to 23%) over the 5’ UTR (6% to 8%) (Figure 4A). RDD preference for 3’ UTR was also observed in deamination-mediated RNA editing (Rosenberg et al., 2011) and in human transcriptome analysis (Li et al., 2011). In mammals, recent findings proposed that 3’ UTRs may play important roles in the regulation of biological processes (Berkovits and Mayr, 2015), probably due to its engagement in intricate RNA base-pairing patterns, which may change in response to protein binding and impact the recruitment of ribosomes particles (Campbell et al., 2012; Castello et al., 2012) as well as, modulate translation (Mayr, 2017) at the 3’ UTR. In plants it is well established that UTRs represent important genetic components controlling gene expression and regulating mRNAs and protein interactions (Hernando et al., 2017; Petrillo et al., 2014). Previous works already suggested that such regulation acts through a wide range of mechanisms that, under diverse cellular conditions, enables a rapid and dynamic response in order to recode gene information (Alvarez et al., 2016; Gallegos and Rose, 2015; Meyer et al., 2015; Yang et al., 2012).

**Figure 4.**
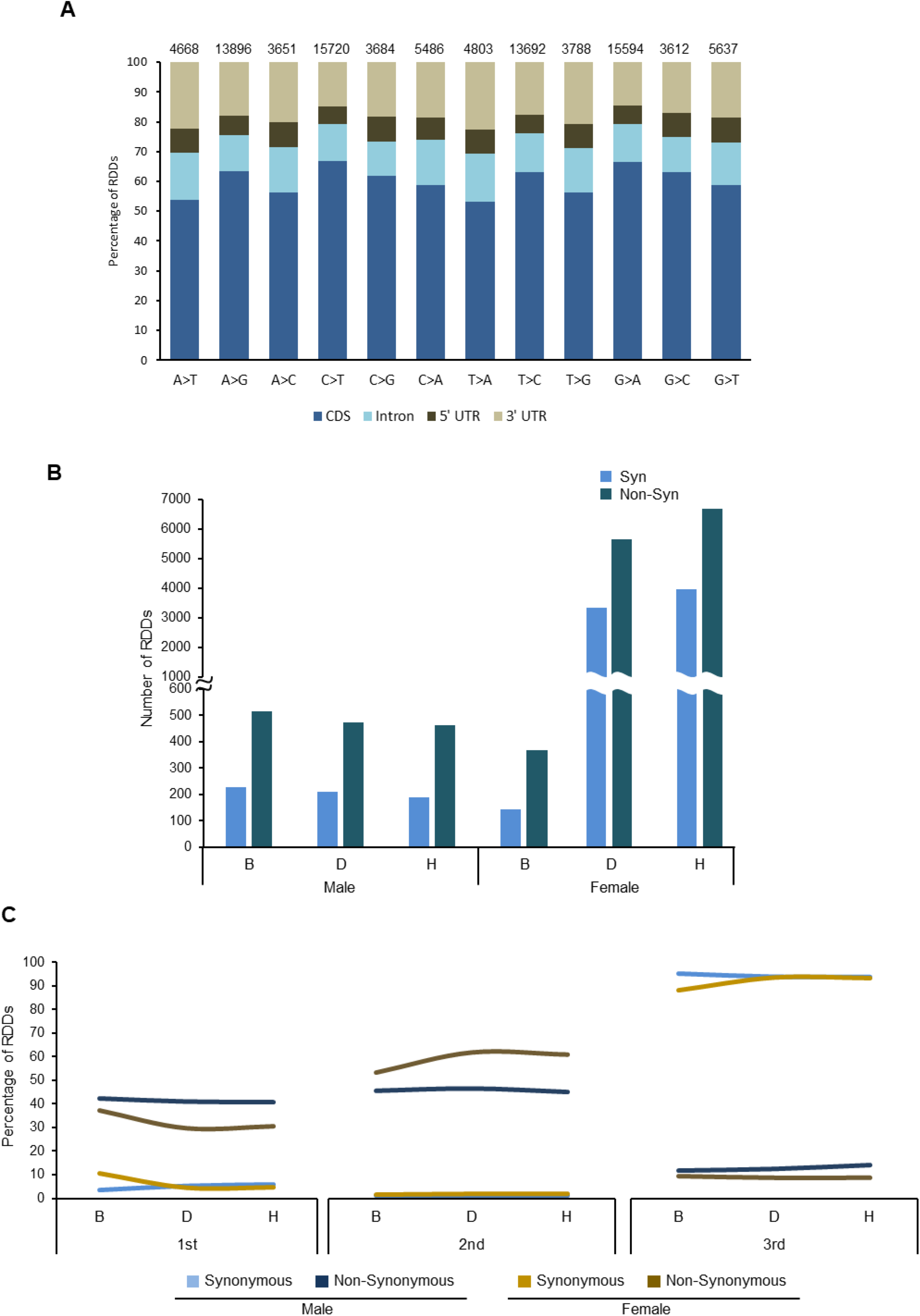
Genomic regions and protein synthesis are differentially affected by RDDs. (A) The combined analysis of both genomes show that editing events are prevalent in 3’ UTR rather than in 5’ UTR. Also, the percentage of RDD present in introns remains roughly the same for all modification, C-to-T being the exception, where a decrease in RDD in the introns is collinear with an increase in RDD events in the CDS regions of transcripts. The numbers above de graphic columns represent the number of editing events. (B) The majority of the identified RDD events in the CDS region leads to non-synonyms modifications, which suggests that the RDD is actively promoting protein modifications. (C) Most codon modifications that occur in 1^st^ or 2^nd^ codon positions lead to non-synonymous modifications, in contrast with editing events in the 3^rd^ codon positions, which lead primarily to synonymous modifications and therefore do not affect the protein that will be translated. See Figure S4 for details.

Further analysis in CDS throughout different inflorescence development stages revealed a greater number of non-synonymous changes, with an average ratio of 2.5 for non-synonymous/synonymous (Figure 4B). A similar pattern was observed in human transcriptome B cells, where 71% of RDDs in coding regions result in non-synonymous amino acid changes (Li et al., 2011). Female inflorescences showed a surprising increase of both modification types in D and H developmental stages, in comparison with the initial B stage (Figure 4B). Non-synonymous/synonymous RDD dynamics in the 1^st^, 2^nd^ and 3^rd^ codon positions were also analyzed. Higher percentages of non-synonymous events in both sexes were observed at the 1^st^ and 2^nd^ codon positions (Figure 4C) being higher for male in the 1^st^ position and for female in the 2^nd^ position. The opposite pattern was observed at the 3^rd^ position where synonymous modifications, as expected by the genetic code, reached the highest value in both sexes (Figure 4C). Exhaustive searches for correlations between the number of editing events/chromosome, number editing events/chromosome length (Figure S3A) or even the number of editing events/gene length (Figure S3B), lead to the conclusion that RDDs distribution is independent of any such parameters (Figure S3).

The fact that RNA editing is widespread among all genomic region and the existence of complex non-synonymous patterns associated to developmental stages, suggests a multiplicity of processes underlying these types of events, probably in accordance with requiring transient synthesis of distinct proteins for each sex.

### Validation of RDDs sites by Sanger sequencing

Since the inflorescence samples, at different developmental stages used in this work, came from the same individuals, we validated the RNA edited sites identified through extensive Sanger sequencing of other plants, distinct from those initially sequenced (female: Fe1_2014 and male: Ma1_2014) and collected in subsequent years (female plants: Fe1_2018, Fe2_2018, Fe3_2015; male plants: Ma1_2018 and Ma2_2018). In order to choose several genes, Integrative Genomic Viewer (IGV) (Robinson et al., 2011) was used. The initially obvious choices were homeotic genes involved in the ABCDE model for flower development (Rijpkema et al., 2010), but none of those transcripts presented nucleotide editing (data not shown). Thus it seems that in *Vitis*, genes considered essential for flowering are rarely an editing target, as was also reported in humans (Xu and Zhang, 2014). Distribution of RDDs in CDS, intron, 5’ UTR and 3’ UTR, was then analyzed in randomly chosen genes of both female and male plants (Table S2). In the majority of the selected genes, the male flowers are heterozygous in the edited sites or do not present RNA editing. Nevertheless, we performed Sanger amplifications for both sexes and the non-edited sequencings were not included in the manuscript. Additionally, we evaluated the *in silico* impact of RDD in the amino acid sequence.

One of the selected genes was *VviSPLAYED* (*VviSYD*) (Kwon et al., 2005; Walley et al., 2008). In female plants, six RDD events were detected by IGV (Figure 5Ai) and validation by Sanger sequencing proved the existence of such six RDD events, only in developmental stage H of female flowers (Figure 5Aii). The first RDD observed was a transition of a thymine to a cytosine (T-to-C) which however, does not lead to a change in the amino acid sequence (Figure 5Aiii). The second RDD event in *VviSYD* was a transversion from a guanine to a thymine (G-to-T) that leads to a change in VviSYD amino acid sequence from valine to leucine (Figure 5Aiii). The third RDD event, was also a transversion, without consequences in the amino acid sequence (Figure 5Aiii). The three remaining RDD were all transversions leading to changes in the amino acid sequence, from aspartic acid to glycine, glycine to valine and valine to serine, respectively (Figure 5Aiii). These results showed that the amino acid sequence of VviSYD protein at female flower development stage H seems to be different than the expected from the coded DNA sequencing. RDD events varying during development were also observed in other genes, such as *VviMYB*-like gene (Figure 5Bi) where a T-to-C transversion leading to a putatively different protein only occurs at stage D, being the edited transcript absent both at the B and H stages of flower development (Figures 5Bi and Bii; Figure S4). On the other hand, the VIT_208s0007g5630 gene, that codes for an uncharacterized protein, presented unedited mRNAs at B stage, although in developmental stages D and H, two distinct transcripts (Figures 5Ci and ii) were detected, due to RNA editing events that can be translated into two different proteins (Figure 5Ciii). Putative somatic mutations, that could had occurred during development, were discarded, by DNA sequencing at male and female flower developmental stages (example for VIT_201s0010g03460 on Figure S5).

**Figure 5.**
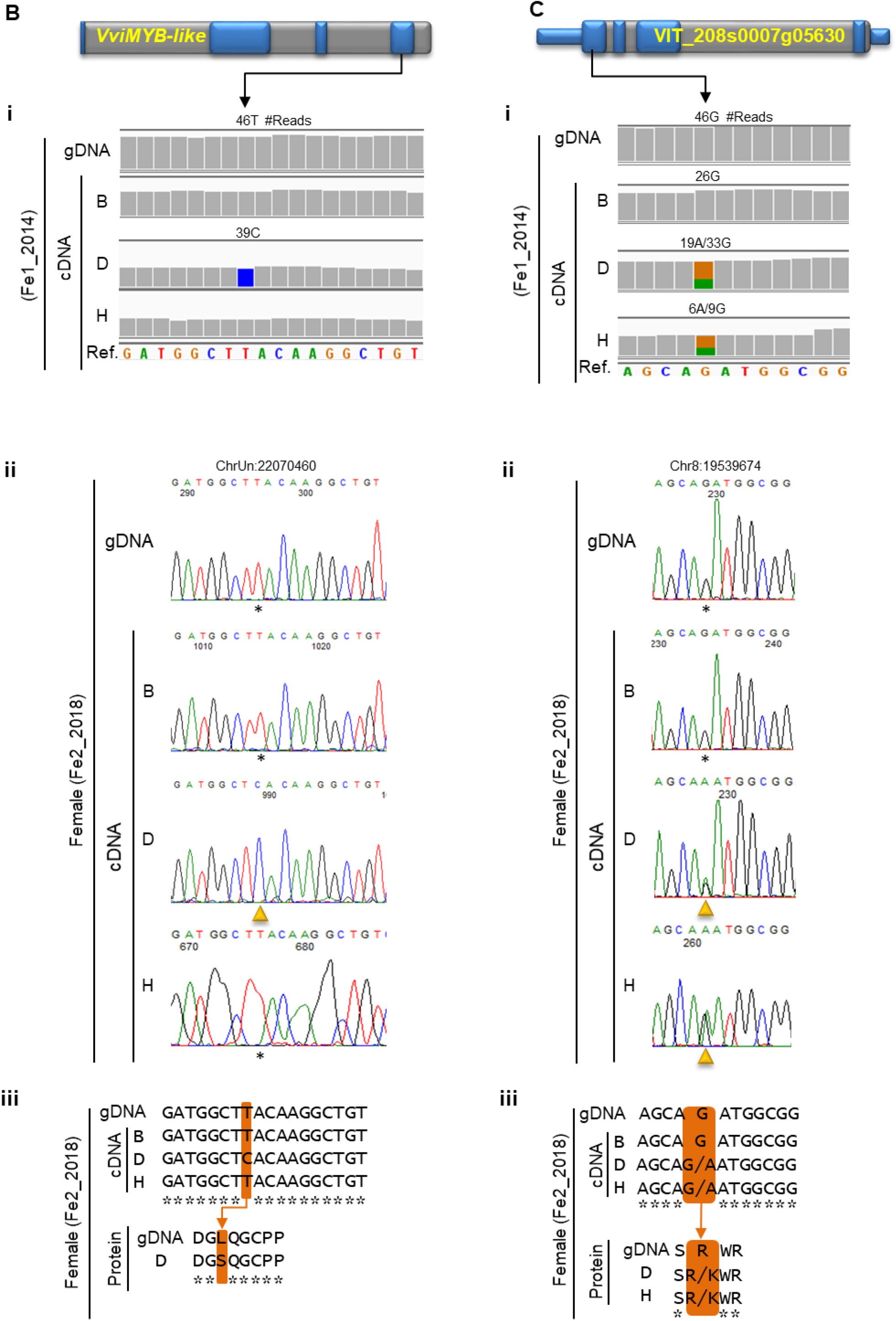
RDD events in *VviSPLAYED*, *VviMYB* and VIT_2085s0007g05630 genes. (A) (i, ii, iii) RDD sites in the *VviSPLAYED* (VIT_205s0020g02000) gene in female inflorescences. (i) *VviSPLAYED* gene structure where blue and grey bars represent exons and introns, respectively. Through the visualization of DNA-seq and RNA-seq six RDD events within the *VviSPLAYED* are observed. (ii) Results were validated, in a second female plant (Fe2_2018), through Sanger sequencing. Electropherogram confirms the six RDD sites at the development stage H and the types of modifications: T-to-C, G-to-T, T-to-A, A-to-G and G-to-C. (iii) *In silico* translation of the edited mRNA, indicates that four (out of six) RDDs are non-synonymous, promoting the emergence of a new protein isoform at stage H. (D) (i, ii, iii) RDD event in the *Vvimyb* (VIT_200s0299g00060) gene in the female inflorescences. (i) *VviMYB* gene structure where blue and grey bars represent exons and introns, respectively. One transient editing event, T-to-C, is observed in developmental stage D. (ii) Validation by Sanger sequencing show the RDD event in a second female plant, Fe2_2018, during developmental stage D. (iii) *In silico* analysis showed this RDD event to be a non-synonymous modification (see Figure 4B and 4C for more details). (G) (i, ii, iii) mRNA two-variants were detected in the VIT_208s0007g055630 gene in the female flowers. (i) VIT_208s0007g055630 gene structure where blue and grey bars represent exons and introns, respectively. The event occurred in D and H developmental stages, with the presence of edited and unedited mRNA. (ii) Validation was performed by Sanger sequencing in a second female plant (Fe2_2018) and mRNA two-variants is observed. (iii) *In silico* analysis showed that the edited mRNA-variant gives rise to a non-synonymous change in protein, from an arginine to a lysine. B, D and H, flower developmental stages; gDNA, genomic DNA.

Altogether, this validation confirms that RNA editing is a nonrandom event, being specific of each flower developmental stage in all individuals of the same sex, analyzed at subsequent years (Figures 5 and S4).

### mRNA and pre-mRNA are targets for editing events

As RNA may be edited at different steps from DNA-to-protein synthesis, we evaluated if RNA editing in *Vitis* occurred before or after RNA splicing. cDNAs were amplified with primers designed for introns flanking exons where editing events were detected. For clarity, we designated as “pre-cDNA” sequences amplified with introns, therefore corresponding to pre-mRNA. Several tests were performed to guarantee that RNA samples were DNA-free (for details see **Methods**). Also, the chosen genes were manually inspected in IGV to intron/exon validation, comparing the number of reads aligning in both regions. In addition, we performed pre-cDNA amplification whenever possible (due to the length of the exon), in the same plant samples used in previous validations.

Each pre-cDNA sequencing was performed independently on at least three replicate samples showing that transcripts from different genes present distinct dynamics: *VviSPLAYED*, is co-transcriptionally edited (Figure 6A) whereas VIT_201s0010g03460, annotated as a methyltransferase, is post-transcriptionally edited (Figure 6B), since the pre-cDNA sequence match with DNA (see Figure S6). These results demonstrate that RNA editing in plants can occur either in pre-mRNA or mRNAs, as was also observed recently in mammalian cells (Mellis et al., 2017). However, we do not know whether the same gene can be both co-transcriptionally and post-transcriptionally edited depending on the needs of the cell/organism.

**Figure 6.**
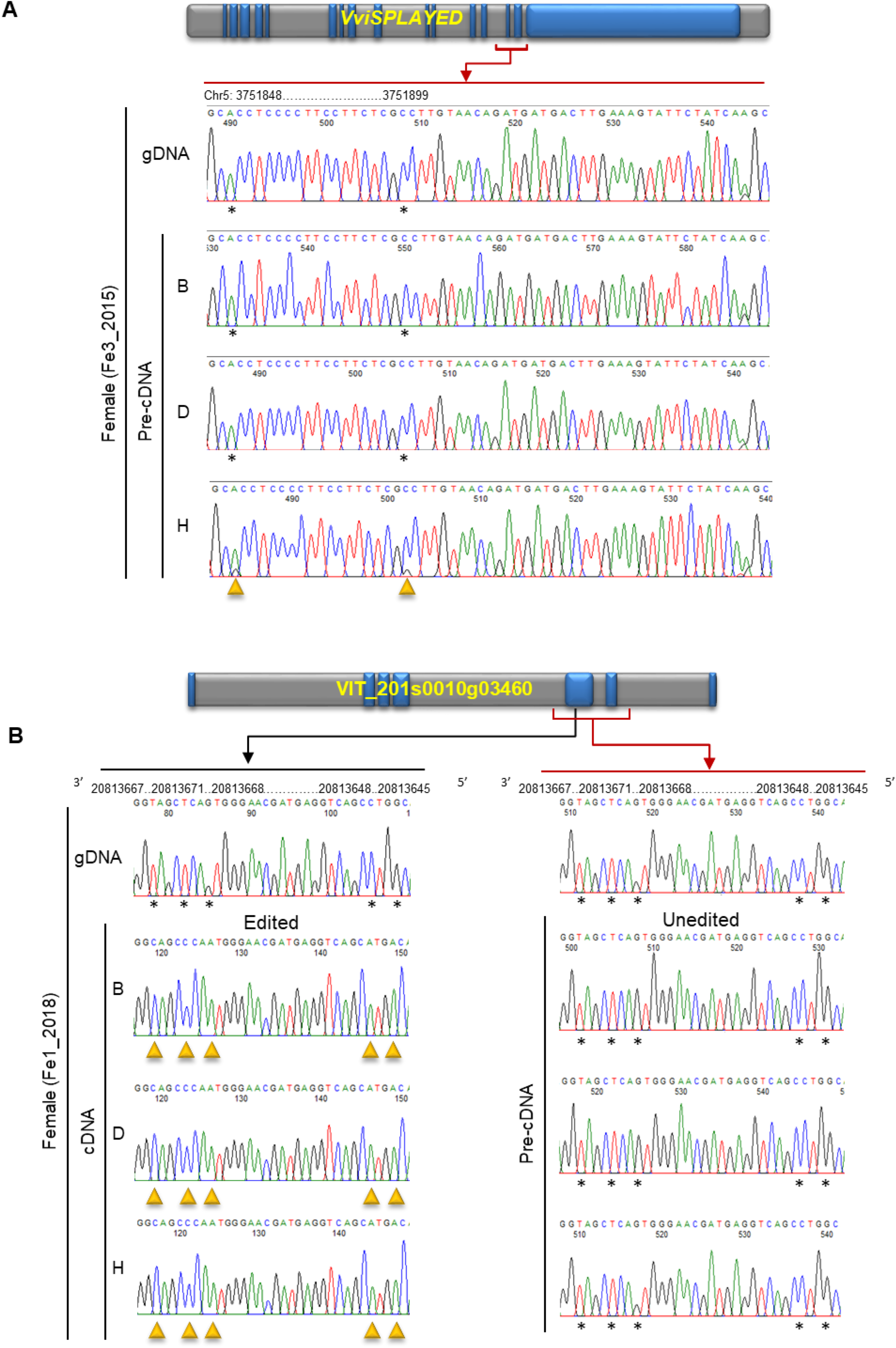
mRNA and pre-mRNA are target for editing events. (A) Amplification with primers for introns that flanked exons allows the determination that *VviSPLAYED* mRNA is co-transcriptionally edited in the last analyzed stage of flower development with two pre-mRNA variants. Previously it has been shown that the transcript of this gene had editing in another exon also in stage H, but with one mRNA variant (see Figure 4A, B and C). (B) We found that post-transcriptionally recoding events could occur in *Vitis* as demonstrated through VIT_201s0010g03460 mRNA that is annotated as a methyltransferase. This transcript shows editing from the beginning of development, but appears unedited when amplified with flaked introns (see Figure S4 and Figure S5).

### Adenosine and cytidine deaminases-like genes are active in *Vitis*

Since the twelve possible types of nucleotide changes were detected in the present work, and that for all of them the mechanism was not yet discovered in plants, we searched for the most well-known editor enzymes responsible for editing A-to-G (adenosine) and C-to-T (cytidine-like), in the *Vitis* genome. These proteins were annotated in plant databases, although no expression studies were until now reported, nor have these enzymes been associated with RNA editing in plants. On *Vitis* chromosome 15, three neighboring genes were found, namely a glucosyltransferase annotated as KDEL motif-containing protein1 (VIT_215s0046g03150) and two genes annotated as adenosine deaminase (VIT_215s0046g03160 and VIT_215s0046g03170). Using the wild grapevine genomes, sequences of these genes in male and female plants were disclosed, reveling a structure similar to the one annotated in the hermaphrodite plant (Figure 7A). To determine if those genes were actively transcribed, we amplified cDNA from the three inflorescence developmental stages, resulting in a unique transcript from the three genes (Figure 7A). The protein alignment revealed conserved motives shared with ADAR in humans (Figure S7A) (Fisher and Beal, 2017).

**Figure 7.**
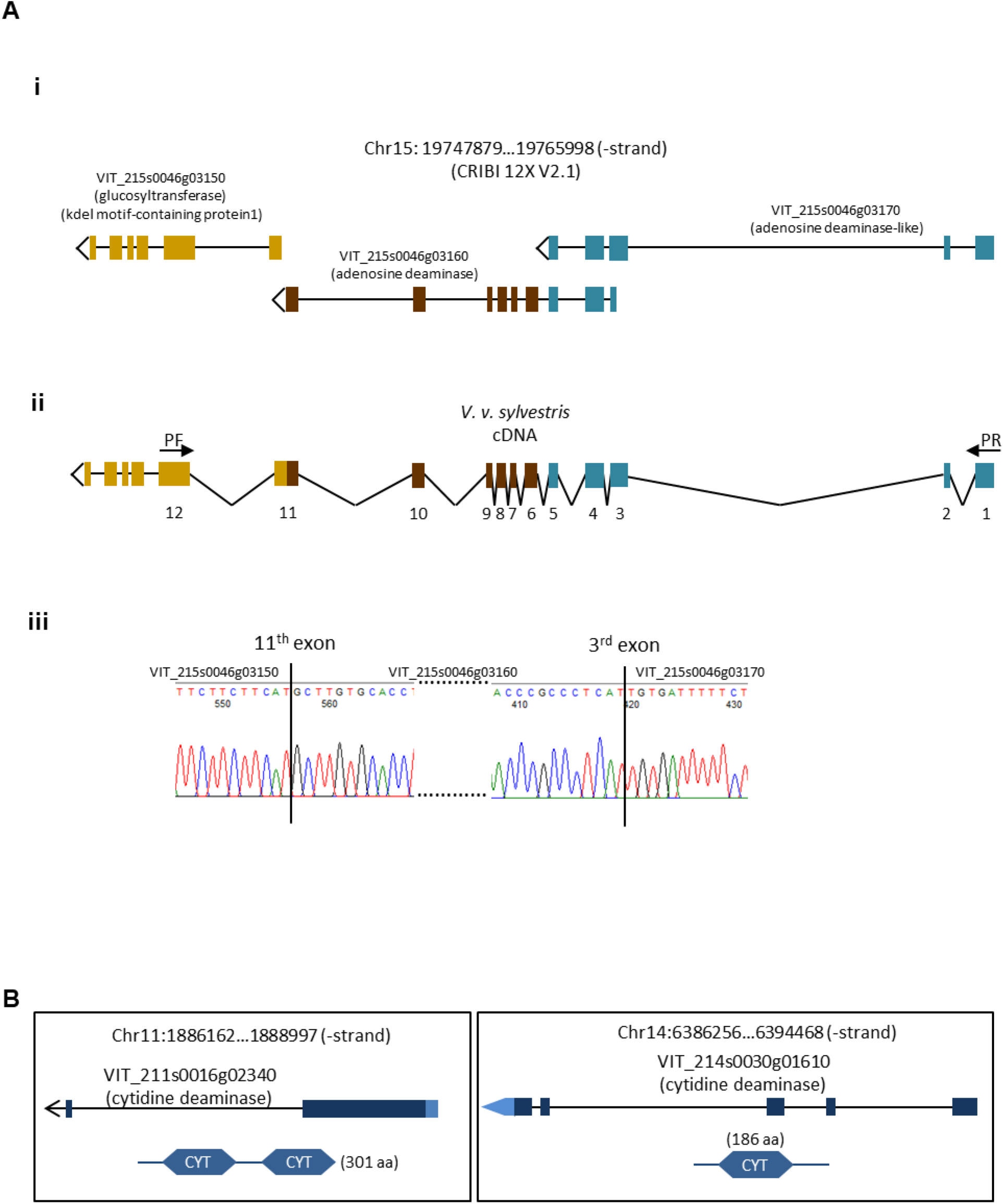
Adenosine and cytidine deaminases-like genes have expression in *Vitis* plants. (A) (i) The *Vitis* genome contains three genes in chromosome 15, two with homology to the adenosine deaminase enzymes (VIT_215s0046g03160 and VIT_215s0046g03170) and a glucosyltransferase gene (VIT_215s0046g03150) annotated as kdel motif-containing protein1. (ii) Amplified in *V. v. sylvestris*, these three genes are not transcribed independently, but as a single mRNA. PF, primer forward; PR, primer reverse. (iii) Sanger sequencing show the continuous transcription of the three annotated genes. (B) Cytidine deaminase-like genes are also present on *Vitis* chromosome 11 and 14 with active transcription. See Figure S7 for details.

We also found two other genes in *Vitis* genome, annotated as cytidine deaminase, one on chromosome 11 (VIT_211s0016g02340) and another one in chromosome 14 (VIT_214s0030g01610) both showing conserved domains namely two and one potential catalytic active deaminase domains, respectively (Figure 7B). In wild *Vitis* both genes are actively transcribed in the three inflorescence development stages analyzed and *in silico* protein alignments revealed a strong homology with other plant proteins (Figure S7B).

Although the real function of these genes was not yet described in any plant species, it is very relevant to report, for the first time, their expression in plants, in opposition to previous affirmations on the inexistence of ADAR-like genes in plants (Jin et al., 2009). Assuming that adenosine and cytidine deaminases have a similar function to the one described for mammals, we propose that ADAR-like gene are responsible for A-to-G (adenosine deaminase) and C-to-T (cytidine deaminase) changes observed in *Vitis*. However, the other ten nucleotide changes remain to be explained.

In other species, mainly in humans, different diseases are associated with altered editing and ADAR activity (Galeano et al., 2013; Han et al., 2015). Therefore, we wonder if in plants, an altered editing pattern can also be associated with responses to biotic or abiotic factors.

Altogether, the present work markedly contributes to deeper understand the process of rewriting genome information and to unveil its differential relevance, accordingly with the sex of the organism in dioecious species evolution.

## DISCUSSION

The first evidence of RNA editing in *Vitis* arose from the comparative analysis of transcriptomes during different stages of inflorescence development (Ramos et al., 2014). Our first assumption was that we were in the presence of SNPs, since we were comparing transcriptomes of two dioecious individuals (wild type) with the annotated genome of a hermaphrodite plant (Ramos et al., 2014; Ramos et al., 2017). After sequencing each of the wild-type genomes and comparing them with their own transcriptomes, it was clear that changes in nucleotides were taking place at post-transcriptional level. As the same plants were analyzed at three distinct flower developmental stages, it is completely excluded the hypothesis that RNA editing is a legacy of a system used to correct RNA transcripts, after harmful DNA mutations (Chen, 2013). We evidence that RNA editing does not persist through the development of inflorescences, being specific of each stage in all individuals of the same sex. These conclusions demonstrate that RNA editing is a highly controlled process regulated during development. Although we did not initially exclude an editing response associated to environment factors, that hypothesis must also be discarded, as the same RNA events are reproduced in subsequent years in the same sites and with the same nucleotide changes. The next big challenge is to unravel the mechanism underling such a fine tuning process, responsible for specific and transient RNA modifications associated to *Vitis* gender.

## Methods

The genome and the transcriptome of two individuals (male and female) from *Vitis vinifera sylvestris* (*V. v. sylvestris*), were compared to determine RNA-DNA differences. The perennial characteristics of *V. v. sylvestris* allowed the study of the same set of plants in different years and developmental stages. This section describes the methods used for these analysis.

### Wild *Vitis* genome sequencing

Fresh leaves from individuals of *V. v. sylvestris* male (Ma1_2016) and female (Fe1_2016) collected in 2016, in the Portuguese Ampelographic Collection, property of the Instituto Nacional de Investigação Agrária e Veterinária, Dois Portos (Lisbon district, Portugal) and were immediately frozen in liquid nitrogen. DNA was extracted using DNeasy Plant Mini Kit (Qiagen). Genomes sequence were obtained on an Illumina Hi-Seq 4000 sequencer, generating 2×125 bp reads for each genome. Bases with low quality (phred quality below 15) were removed from the 3’ and 5’ ends and only mate pairs (forward and reverse reads) were used for the next analysis. Reads were then mapped to the reference genome (PN40024 - 12X; CRIBI, Grape genome database, hereafter as reference genome), using BWA (Li and Durbin, 2009) with the default parameters. Assemblies were optimized using Picard to remove PCR duplicates (http://picard.sourceforge.net). SNP calling was performed using GATK’s Haplotype Caller (DePristo et al., 2011; McKenna et al., 2010) and SNP context was annotated by snpEff (http://snpeff.sourceforge.net). Filters set to the SNP calling were SnpCluster (more than 2 SNPs found in cluster of size=10), LowQual (SNP with quality score < 30), LowCov (SNP with coverage < 20) and Mask (SNP is at least 10 base near to indel location). For more details, refer to Ramos et al., 2020.

### Wild *Vitis* transcriptomic analysis

RNA was obtained from three inflorescence developmental stages (B, D and H), from the same individuals used in genome sequencing (Ma1_2014 and Fe1_2014) (Ramos et al., 2014). Total RNA was extracted with Spectrum Total RNA kit (Sigma-Aldrich, Inc, Spain), according to the manufacturer’s instructions. Poly-A tails were used to cDNA library synthesis along Illumina standard pipeline. The obtained transcriptomes were assembled against the reference public genome (PN40024 - 12X V1) (Vitulo et al., 2014). A variant caller software was used to determine SNP between the sequenced transcriptomes and the reference genome. Calls were discarded if: (1) the SNP call was within the first or last 60 bases of a linkage group sequence; (2) the reference genome base was ambiguous at the SNP position or within 60 bases of it; (3) another SNP was detected within 50 bases; (4) the average copy number of the SNP flanking region was more than two; (5) the Illumina quality score of either allele was less than 20 and (6) the number of reads supporting either allele was less than two reads per accession.

The initial list of variants was filtered using the Phred quality scores of the position and surrounding bases. To reduce false positives in variant calling the minimum variant frequency was set to 15%. The minimum coverage required was 4 reads. Gene expression was measured using the number of Reads per Kilobase of exon model per Million mapped reads (RPKM).

### Computational strategy for RNA-DNA editing sites analysis

The comparison between DNA and RNA, obtained from the Next Sequencing Generation, was performed using a RStudio (1.0.143) (Team, 2016) cluster, running R (3.4.3) (Team, 2017). In this analysis, the genome from each individual (Ma1 and Fe1) was compared to the respective three transcriptomes, corresponding to different inflorescence development stages, in order to detect sites with RNA-DNA differences (RNA editing) across the development stages. An in-house R script was developed to determine RNA editing sites with the following criteria: ≥ 5 reads covering each putative event site and RPKM ≥ 1 at the six samples (male and female). Events that could compromise the analysis, such as deletions, inserts and indels have been removed. The applied filters in genome sequencing of *V. v. sylvestris* (referred as DNA-seq) and RNA-seq led to a situation where information was missing between DNA-seq/reference or between RNA-seq/reference and these cases required a deeper analysis (Figure 1).

Situations without information regarding the number of Adenosines (A), Cytosines (C), Guanines (G) and Thymines (T) covering the *loci* were interpreted as a consequence of: (1) the sequenced nucleotides on DNA-seq/RNA-seq were identical to the reference genome, or (2) the coverage was insufficient to be considered. To avoid situations of false editing, on such of those situations the number of occurrences of each base covering those specific *loci* were counted. This approach provided information regarding the number of As, Cs, Gs and Ts covering all situations that were lacking due to the initial filters (described above).

The developed in-house script, to compare RNA and DNA, was based in the following four main steps: (1) Nucleotides transcribed from the anti-sense strand of the reference genome, were switched to its complementary base (in order to make them comparable with the genome) and the chromosomal positions were calculated to all variants; (2) To determine if a genomic site was homozygous or heterozygous we considered the frequency (*freq*) of readings for each nucleotide based on the following criteria: *freq* ≥ 0.90, homozygous; *freq* < 0.90, heterozygous. If any observed nucleotide was found with a *freq* < 0.10, it was considered a sequencing error and discarded; (3) For RNA-seq, it was applied the same criteria used to infer the number of mRNA variants; and (4) Cases where the observed RNA-seq nucleotide was not found in the DNA-seq, were considered RNA editing events. The global analysis was performed with both homozygous and heterozygous gDNA sites (Figure 2). Subsequently, RNA editing was studied only in homozygous sites.

To discard a possible enrichment of chromosome editing, we analyzed the distribution of editing events in male and female plants, calculating the ratio between the number of RDDs and the length of the chromosome (Figure S3).

All in-house scripts are available upon request.

### Refining knowledge on RNA editing sites

Some RNA editing events detected by the in-house R scripts were manually inspected using the Integrated Genome Viewer (IGV) browser (Brooks et al., 2014; Manavalan et al., 2012; Xiong et al., 2005), loaded with the reference genome (PN40024 – 12X), the most recent gene annotation (V2.1) and the Binary Alignment Map (BAM) files corresponding to the genome and transcriptomes sequencings.

The RNA editing events detected were deeply inspected. The genomic environment of each RNA editing site was evaluated considering gene annotation (V2.1) discarding multiple transcripts, with different contexts, covered the RNA editing site (Figures 2E and 4A).

For each of the twelve modifications discovered, the frequency of each nucleotide up to 1000 nt upstream and downstream the RNA editing site was calculated. No major differences were observed in the overall analysis, except in the nucleotide immediately preceding or following some the RNA editing events. For the propose of this manuscript, only the ± 10 nt are shown (Figure 3C).

### Plant material for RNA editing validation

Inflorescences from female and male plants of *V. v. sylvestris* were collected in Portuguese Ampelographic Collection (PRT051), property of the Instituto Nacional de Investigação Agrária e Veterinária, Dois Portos, Lisbon, Portugal. The harvesting of inflorescence/floral buds was individually made from several plants of each flower type (male and female), by the end of April and the beginning of May. Inflorescences were classified according to its phenological developmental stage as B, D and H (Baggiolini, 1952) (Figure 1). Since this work was based on phenological stages, samples were always collected by the same operator at the same time-frame between each sampling, in order to guarantee its accuracy. To study RNA editing dynamic across the years, samples were collected between 2014 and 2018 from the same individuals used on the Next Generation Sequencing, already described, as well as, from other wild female (Fe1_2018, Fe2_2018, Fe3_2015) and male (Ma1_2018 and Ma2_2018) plants.

### RNA and DNA extraction

Inflorescence buds, at different developmental stages, were individually reduced to powder using liquid nitrogen. To exclude putative mutations that may occur along the development of the new branches, total RNA and DNA were extracted from the same powder, using Spectrum™ Plant Total RNA Kit (Sigma-Aldrich) and DNeasy® Plant Mini Kit (Qiagen, Valencia, CA, USA), respectively, following manufacturer’s instructions. Total RNA was additionally treated with 30 units of DNase I (Qiagen) and an intron-spanning PCR product was checked on an agarose gel to exclude the possibility of DNA contamination. Nucleic acid concentration was measured using a microplate reader Synergy HT (Biotek, Germany), with the software Gen5™ (Biotek, Germany).

### cDNA synthesis

cDNA and pre-cDNA were synthesized from 100 ng of total RNA, using oligo(dT)_20_ and RevertAid Reverse Transcriptase (Thermo Fisher Scientific™) in a 20 μL-reaction volume, following the manufacturer’s instructions. cDNA concentration was measured as described above.

### Gene selection, amplification and Sanger sequencing

Genes to validate the global analysis were first chosen by having multiple editing events, manually inspected using IGV (Integrative Genomic Viewer) (Fernandez et al., 2007; Sreekantan et al., 2006; Xiong et al., 2005), loaded with the reference genome (12X), the V2.1 annotation (Vitulo et al., 2014) and the BAM files obtained from male and female genomes and RNA-seq. Primers were designed using Primer Premier5 considering a primer length of 20 ± 2 bp and melting temperature of *circa* 55ºC (Table S2).

The DNA and the corresponding cDNA (synthesized from RNA extracted from the same powder as DNA) amplifications were performed through PCR in 50 μL total volume composed by 2 µg of DNA (or cDNA), PCR buffer (20 mM Tris-HCl [pH 8.4], 50 mM KCl), 1.5 mM of Mg_2_, 0.2 mM of dNTP mix, 0.4 μM of each forward and reverse primers, 10 U of Taq DNA polymerase and autoclaved MiliQ water. The applied program had 3 min for initial denaturation at 95°C followed by 34 cycles of 45 s at 95°C (denaturation), 45 s at 55°C (annealing) and 90 s at 72°C (extension), followed by a final extension for 5 min at 72°C. For amplification of the pre-mRNA both primers were designed for the intron that flanks the exon were it was previously detected RNA editing (Table S2).

PCR amplifications were inspected on an agarose gel (1.7%). The remaining product was purified using PureLink™ Quick PCR Purification kit (Thermo Fisher Scientific™), according to the kit protocol. For each gene DNA and cDNA were Sanger sequenced in three technical replicates without resorting to cloning. The provided electropherograms were manually inspected.

## AUTHOR CONTRIBUTIONS

M.R. and M.M.R.C. conceived the project and were responsible for project initiation. M.R., M.J.N.R. and J.L.C. designed the experiments. M.R. supervised and managed the project and research. M.J.N.R. and D.F-S. designed and performed the in-house scripts for bioinformatic analysis. M.J.N.R., D.F-S., H.S. and M.R. performed the experiments. M.J.N.R., D.F-S., H.S. and M.R. analyzed the data. J.C., M.R., M.J.N.R. and J.L.C. established *Vitis vinifera sylvestris* collection and collected plant tissues according phenological developmental stages. The manuscript was organized and written by M.J.N.R., J.L.C. and M.R. All authors discussed the results and commented on the manuscript. Correspondence and requests for materials should be addressed to M.R. or M.J.N.R.

## ACKNOWLEDGMENTS

This work was financed by project PTDC/AGR-GPL/119298/2010 from Fundação para a Ciência e Tecnologia (FCT, Portugal) and developed in LEAF (UID/AGR/04129/2013) and BioISI (UID/MULTI/04046/2013), research centres, with the support of Research Infrastructure PORBIOTA (POCI-01-0145-FEDER-022127). Authors J.L.C., M.J.N.R., M.M.R.C. and M.R. were funded by FCT fellowships, respectively, SFRH/BD/85824/2012, SFRH/BD/110274/2015, SFRH/BSAB/113781/2015, SFRH/BPD/64905/2009. We are also grateful to Eng. Eiras-Dias, curator of the Portuguese Ampelographic Collection, Instituto Nacional de Investigação Agrária e Veterinária, Dois Portos for the collaboration in this work allowing the access to the *Vitis* collection. To Mariana G. Lopes and Pedro M. Silvestre our thanks goes for their help in the laboratory. We also want to thanks Prof. Helena Oliveira for the support given to genomes resequencing.

## REFERENCES

Alvarez, D., Voss, B., Maass, D., Wust, F., Schaub, P., Beyer, P., and Welsch, R. (2016). Carotenogenesis is regulated by 5’UTR-mediated translation of phytoene synthase splice variants. Plant Physiol. 172, 2314–2326.

Badouin, H., Velt, A., Gindraud, F., Flutre, T., Dumas, V., Vautrin, S., Marande, W., Corbi, J., Sallet, E., Ganofsky, J., et al. (2020). The wild grape genome sequence provides insights into the transition from dioecy to hermaphroditism during grape domestication. Preprint at https://www.biorxiv.org/content/10.1101/2020.01.07.897082v2.

Baggiolini, M. (1952). Les stades repères dans le developpement annuel de la vigne et leur utilisation pratique. Rev. Romande Agric. Vitic. Arbor. 8, 4–6.

Bahn, J.H., Lee, J.H., Li, G., Greer, C., Peng, G., and Xiao, X. (2012). Accurate identification of A-to-I RNA editing in human by transcriptome sequencing. Genome Res. 22, 142–150.

Bailey, T.L., Boden, M., Buske, F.A., Frith, M., Grant, C.E., Clementi, L., Ren, J., Li, W.W., and Noble, W.S. (2009). MEME SUITE: tools for motif discovery and searching. Nucleic Acids Res. 37, W202–208.

Berkovits, B.D., and Mayr, C. (2015). Alternative 3’ UTRs act as scaffolds to regulate membrane protein localization. Nature 522, 363–367.

Brooks, C., Nekrasov, V., Lippman, Z.B., and Van Eck, J. (2014). Efficient gene editing in tomato in the first generation using the clustered regularly interspaced short palindromic repeats/CRISPR-Associated9 system. Plant Physiol. 166, 1292–1297.

Campbell, Z.T., Bhimsaria, D., Valley, C.T., Rodriguez-Martinez, J.A., Menichelli, E., Williamson, J.R., Ansari, A.Z., and Wickens, M. (2012). Cooperativity in RNA-protein interactions: global analysis of RNA binding specificity. Cell Rep. 1, 570–581.

Castello, A., Fischer, B., Eichelbaum, K., Horos, R., Beckmann, B.M., Strein, C., Davey, N.E., Humphreys, D.T., Preiss, T., Steinmetz, L.M., et al. (2012). Insights into RNA biology from an atlas of mammalian mRNA-binding proteins. Cell 149, 1393–1406.

Chen, L. (2013). Characterization and comparison of human nuclear and cytosolic editomes. Proc. Natl. Acad. Sci. USA 110, E2741–E2747.

Cheng, S., Gutmann, B., Zhong, X., Ye, Y., Fisher, M.F., Bai, F., Castleden, I., Song, Y., Song, B., Huang, J., et al. (2016). Redefining the structural motifs that determine RNA binding and RNA editing by pentatricopeptide repeat proteins in land plants. Plant J. 85, 532–547.

Coito, J.L., Ramos, M., Cunha, J.M.M., Silva, H.G., Amâncio, S., Costa, M.M.R., and Rocheta, M.P. (2017). *VviAPRT3* and *VviFSEX*: two genes involved in sex specification able to distinguish different flower types in *Vitis*. Front. Plant Sci. 8, 98.

DePristo, M.A., Banks, E., Poplin, R., Garimella, K.V., Maguire, J.R., Hartl, C., Philippakis, A.A., del Angel, G., Rivas, M.A., Hanna, M., et al. (2011). A framework for variation discovery and genotyping using next-generation DNA sequencing data. Nat. Genet. 43, 491–498.

Fernandez, L., Torregrosa, L., Terrier, N., Sreekantan, L., Grimplet, J., Davies, C., Thomas, M.R., Romieu, C., and Ageorges, A. (2007). Identification of genes associated with flesh morphogenesis during grapevine fruit development. Plant Mol. Biol. 63, 307–323.

Fisher, A.J., and Beal, P.A. (2017). Effects of Aicardi-Goutieres syndrome mutations predicted from ADAR-RNA structures. RNA Biol. 14, 164–170.

Galeano, F., Rossetti, C., Tomaselli, S., Cifaldi, L., Lezzerini, M., Pezzullo, M., Boldrini, R., Massimi, L., Di Rocco, C.M., Locatelli, F., et al. (2013). ADAR2-editing activity inhibits glioblastoma growth through the modulation of the CDC14B/Skp2/p21/p27 axis. Oncogene 32, 998–1009.

Gallegos, J.E., and Rose, A.B. (2015). The enduring mystery of intron-mediated enhancement. Plant Sci. 237, 8–15.

Glenn, T.C. (2011). Field guide to next-generation DNA sequencers. Mol. Ecol. Resour. 11, 759–769.

Han, L., Diao, L., Yu, S., Xu, X., Li, J., Zhang, R., Yang, Y., Werner, H.M.J., Eterovic, A.K., Yuan, Y., et al. (2015). The genomic landscape and clinical relevance of A-to-I RNA editing in human cancers. Cancer Cell 28, 515–528.

Hernando, C.E., Romanowski, A., and Yanovsky, M.J. (2017). Transcriptional and post-transcriptional control of the plant circadian gene regulatory network. Biochim. Biophys. Acta Gene Regul. Mech. 1860, 84–94.

Ichinose, M., and Sugita, M. (2017). RNA editing and its molecular mechanism in plant organelles. Genes 8, 1–15.

Jaillon, O., Aury, J.M., Noel, B., Policriti, A., Clepet, C., Casagrande, A., Choisne, N., Aubourg, S., Vitulo, N., Jubin, C., et al. (2007). The grapevine genome sequence suggests ancestral hexaploidization in major angiosperm phyla. Nature 449, 463–467.

Jin, Y., Zhang, W., and Li, Q. (2009). Origins and evolution of ADAR-mediated RNA editing. IUBMB Life 61, 572–578.

Kwon, C.S., Chen, C.B., and Wagner, D. (2005). *WUSCHEL* is a primary target for transcriptional regulation by *SPLAYED* in dynamic control of stem cell fate in *Arabidopsis*. Genes and Dev. 19, 992–1003.

Li, H., and Durbin, R. (2009). Fast and accurate short read alignment with Burrows-Wheeler transform. Bioinformatics 25, 1754–1760.

Li, M., Wang, I.X., Li, Y., Bruzel, A., Richards, A.L., Toung, J.M., and Cheung, V.G. (2011). Widespread RNA and DNA sequence differences in the human transcriptome. Science 333, 53–58.

Liscovitch-Brauer, N., Alon, S., Porath, H.T., Elstein, B., Unger, R., Ziv, T., Admon, A., Levanon, E.Y., Rosenthal, J.J.C., and Eisenberg, E. (2017). Trade-off between transcriptome plasticity and genome evolution in cephalopods. Cell 169, 191–202.

Maas, S., Patt, S., Schrey, M., and Rich, A. (2001). Underediting of glutamate receptor GluR-B mRNA in malignant gliomas. Proc. Natl. Acad. Sci. USA 98, 14687–14692.

Manavalan, L.P., Chen, X., Clarke, J., Salmeron, J., and Nguyen, H.T. (2012). RNAi-mediated disruption of squalene synthase improves drought tolerance and yield in rice. J. Exp. Bot. 63, 163–175.

Massonnet, M., Cochetel, N., Minio, A., Vondras, A.M., Muyle, A., Lin, J., Garcia, J.F., Zhou, Y., Delledonne, M., Riaz, S., et al. (2020). The genetic basis of sex determination in grapevines (Vitis spp.). Preprint at https://www.biorxiv.org/content/10.1101/2019.12.11.861377v1.

Mayr, C. (2017). Regulation by 3’-untranslated regions. Annu. Rev. Genet. 51, 171–194.

McKenna, A., Hanna, M., Banks, E., Sivachenko, A., Cibulskis, K., Kernytsky, A., Garimella, K., Altshuler, D., Gabriel, S., Daly, M., et al. (2010). The genome analysis toolkit: a MapReduce framework for analyzing next-generation DNA sequencing data. Genome Res. 20, 1297–1303.

Mellis, I.A., Gupte, R., Raj, A., and Rouhanifard, S.H. (2017). Visualizing adenosine-to-inosine RNA editing in single mammalian cells. Nat. Meth. 14, 801–804.

Meng, Y., Chen, D., Jin, Y., Mao, C., Wu, P., and Chen, M. (2010). RNA editing of nuclear transcripts in *Arabidopsis thaliana*. BMC Genomics 11 Suppl 4, S12.

Meyer, K.D., Patil, D.P., Zhou, J., Zinoviev, A., Skabkin, M.A., Elemento, O., Pestova, T.V., Qian, S.B., and Jaffrey, S.R. (2015). 5’ UTR m(6)A promotes cap-independent translation. Cell 163, 999–1010.

Nishikura, K. (2006). Editor meets silencer: crosstalk between RNA editing and RNA interference. Nat. Rev. Mol. Cell Biol. 7, 919–931.

Okuda, K., Chateigner-Boutin, A.L., Nakamura, T., Delannoy, E., Sugita, M., Myouga, F., Motohashi, R., Shinozaki, K., Small, I., and Shikanaia, T. (2009). Pentatricopeptide repeat proteins with the DYW motif have distinct molecular functions in RNA editing and RNA cleavage in *Arabidopsis* chloroplasts. Plant Cell 21, 146–156.

Okuda, K., Myouga, F., Motohashi, R., Shinozaki, K., and Shikanai, T. (2007). Conserved domain structure of pentatricopeptide repeat proteins involved in chloroplast RNA editing. Proc. Natl. Acad. Sci. USA 104, 8178–8183.

Petrillo, E., Godoy Herz, M.A., Barta, A., Kalyna, M., and Kornblihtt, A.R. (2014). RNA Biol. RNA Biol. 11, 1215–1220.

Picq, S., Santoni, S., Lacombe, T., Latreille, M., Weber, A., Ardisson, M., Ivorra, S., Maghradze, D., Arroyo-Garcia, R., Chatelet, P., et al. (2014). A small XY chromosomal region explains sex determination in wild dioecious *V. vinifera* and the reversal to hermaphroditism in domesticated grapevines. BMC Plant Biol. 14, 229.

Qulsum, U., Azad, M.T.A., and Tsukahara, T. (2019). Analysis of tissue-specific RNA editing events of genes involved in RNA editing in *Arabidopsis thaliana*. J. Plant Biol. 62, 351–358.

Ramos, M.J., Coito, J.L., Silva, H.G., Cunha, J., Costa, M.M., and Rocheta, M. (2014). Flower development and sex specification in wild grapevine. BMC Genomics 15, 1095.

Ramos, M.J.N., Coito, J., Fino, J., Cunha, J., Silva, H., de Almeida, P.G., Costa, M.M.R., Amancio, S., Paulo, O.S., and Rocheta, M. (2017). Deep analysis of wild *Vitis* flower transcriptome reveals unexplored genome regions associated with sex specification. Plant Mol. Biol. 93, 151–170.

Ramos, M.J.N., Coito, J.L., Silva, D.F., Cunha, J., Costa, M.M.R., Amâncio, S., and Rocheta, M. (2020). Portuguese wild grapevine genome re-sequencing. Preprint at https://doi.org/10.1101/2020.02.19.955781.

Rijpkema, A.S., Vandenbussche, M., Koes, R., Heijmans, K., and Gerats, T. (2010). Variations on a theme: Changes in the floral ABCs in angiosperms. Semin. Cell Dev. Biol. 21, 100–107.

Robinson, J.T., Thorvaldsdottir, H., Winckler, W., Guttman, M., Lander, E.S., Getz, G., and Mesirov, J.P. (2011). Integrative genomics viewer. Nat. Biotechnol. 29, 24–26.

Rosenberg, B.R., Hamilton, C.E., Mwangi, M.M., Dewell, S., and Papavasiliou, F.N. (2011). Transcriptome-wide sequencing reveals numerous APOBEC1 mRNA-editing targets in transcript 3’ UTRs. Nat. Struct. Mol. Biol. 18, 230–236.

Rudinger, M., Polsakiewicz, M., and Knoop, V. (2008). Organellar RNA editing and plant-specific extensions of pentatricopeptide repeat proteins in jungermanniid but not in marchantiid liverworts. Mol. Biol. Evol. 25, 1405–1414.

Salter, J.D., Bennett, R.P., and Smith, H.C. (2016). The APOBEC protein family: united by structure, divergent in function. Trends Biochem. Sci. 41, 578–594.

Schirmer, M., D’Amore, R., Ijaz, U.Z., Hall, N., and Quince, C. (2016). Illumina error profiles: resolving fine-scale variation in metagenomic sequencing data. BMC Bioinformatics 17.

Sreekantan, L., Torregrosa, L., Fernandez, L., and Thomas, M.R. (2006). VvMADS9, a class B MADS-box gene involved in grapevine flowering, shows different expression patterns in mutants with abnormal petal and stamen structures. Funct. Plant Biol. 33, 877–886.

Team, R. (2016). RStudio: Integrated Development for R. RStudio, Inc., Boston, MA URL http://www.rstudio.com/.

Team, R.C. (2017). R: A language and environment for statistical computing. R Foundation for Statistical Computing, Vienna, Austria. Available online at https://www.R-project.org/.

Vitulo, N., Forcato, C., Carpinelli, E.C., Telatin, A., Campagna, D., D’Angelo, M., Zimbello, R., Corso, M., Vannozzi, A., Bonghi, C., et al. (2014). A deep survey of alternative splicing in grape reveals changes in the splicing machinery related to tissue, stress condition and genotype. BMC Plant Biol. 14, 1–16.

Walley, J.W., Rowe, H.C., Xiao, Y., Chehab, E.W., Kliebenstein, D.J., Wagner, D., and Dehesh, K. (2008). The chromatin remodeler *SPLAYED* regulates specific stress signaling pathways. PLoS Patho. 4, e1000237.

Wang, I.X., So, E., Devlin, J.L., Zhao, Y., Wu, M., and Cheung, V.G. (2013). ADAR regulates RNA editing, transcript stability, and gene expression. Cell Rep. 5, 849–860.

Xiong, A.S., Yao, Q.H., Peng, R.H., Li, X., Han, P.L., and Fan, H.Q. (2005). Different effects on ACC oxidase gene silencing triggered by RNA interference in transgenic tomato. Plant Cell Rep. 23, 639–646.

Xu, G., and Zhang, J. (2014). Human coding RNA editing is generally nonadaptive. Proc. Natl. Acad. Sci. USA 111, 3769–3774.

Yang, X., Zhang, H., and Li, L. (2012). Alternative mRNA processing increases the complexity of microRNA-based gene regulation in *Arabidopsis*. Plant J. 70, 421–431.

Zhou, Y.F., Massonnet, M., Sanjak, J.S., Cantu, D., and Gaut, B.S. (2017). Evolutionary genomics of grape (*Vitis vinifera* ssp. *vinifera*) domestication. Proc. Natl. Acad. Sci. USA 114, 11715–11720.

